# Characterizing Building Blocks of Resource Constrained Biological Networks

**DOI:** 10.1101/352450

**Authors:** Yuanfang Ren, Ahmet Ay, Alin Dobra, Tamer Kahveci

## Abstract

Identification of motifs-recurrent and statistically significant patterns-in biological networks is the key to understand the design principles, and to infer governing mechanisms of biological systems. This, however, is a computationally challenging task. This task is further complicated as biological interactions depend on limited resources, i.e., a reaction takes place if the reactant molecule concentrations are above a certain threshold level. This biochemical property implies that network edges can participate in a limited number of motifs simultaneously. Existing motif counting methods ignore this problem. This simplification often leads to inaccurate motif counts (over-or under-estimates), and thus, wrong biological interpretations. In this paper, we develop a novel motif counting algorithm, *Partially Overlapping MOtif Counting* (*POMOC*), that considers capacity levels for all interactions in counting motifs. Our experiments on real and synthetic networks demonstrate that motif count using the *POMOC* method significantly differs from the existing motif counting approaches, and our method extends to large-scale biological networks in practical time. Our results also show that our method makes it possible to characterize the impact of different stress factors on cell’s organization of network. In this regard, analysis of a *S. cerevisiae* transcriptional regulatory network using our method shows that oxidative stress is more disruptive to organization and abundance of motifs in this network than mutations of individual genes. Our analysis also suggests that by focusing on the edges that lead to variation in motif counts, our method can be used to find important genes, and to reveal subtle topological and functional differences of the biological networks under different cell states.

---

**Figure 1:**
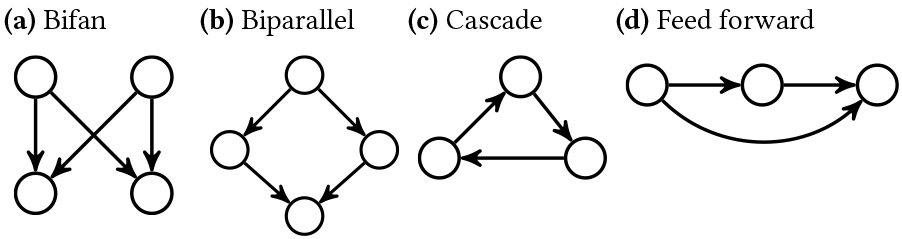
Four conserved motifs studied frequently in the literature.

## 1 INTRODUCTION

Biological networks describe interactions between molecules such as genes and proteins [2, 6]. These networks are often modeled as graphs where nodes and edges represent molecules and their interactions, respectively. Biological networks are involved in many key biological processes including transcriptional regulation, interactions between a cell and its environment, and controlling a cell’s specificity [8, 27, 28]. Understanding biological networks is essential for understanding how cells function. Efforts on computational analysis of biological networks have been growing rapidly in recent years as large-scale data collection at low cost is now possible.

One of the most fundamental challenges in computational network studies is the motif counting problem. A network motif is a pattern of local interconnections (i.e., a small subnetwork) observed significantly more frequently in a given network than in a random network of the same size [3, 32]. Existing studies have already uncovered existence of network motifs such as feed forward loop and bifan (see Figure 1) [23, 31]. Motifs utilize the basic control mechanisms to govern biologically important dynamic behaviors, such as oscillations, generation of molecular pulses, and rapid or delayed responses [3, 37]. Thus, the presence or relative abundance of motifs in biological networks is often used to characterize their topology, function, and robustness [16, 29]. Network motifs have been effectively used to study the biological processes that regulate transcription [31], to find the genetic factors that impact various diseases [17, 21] and to discover new drugs [33].

Identifying motifs and counting them in biological networks is a computationally challenging task as it requires solving the subgraph isomorphism problem, which is NP-complete [15]. Several methods have been developed to count instances of a motif in a given network [1, 18, 19, 30]. These methods could be categorized into two classes [18]. The first class counts all instances of a given motif ignoring the fact that some motifs may share edges (*F*_1_ measure) [30]. The second class counts all non-overlapping instances of a given motif, i.e., those which do not share any edge (*F*_2_ measure). Figure 2 shows the difference between these two frequency measures on a hypothetical network *G*. Consider the motif pattern *M* in Figure 2b. Our input network *G* in Figure 2a yields six possible embeddings of *M* shown in Figures 2d-2i. Thus, *F*_1_ measure of *M* in *G* is six. However, out of these six embeddings at most two can be chosen without picking the same edge multiple times (e.g., Figures 2d and 2i). Thus, *F*_2_ measure of *M* in *G* is two.

**Figure 2:**
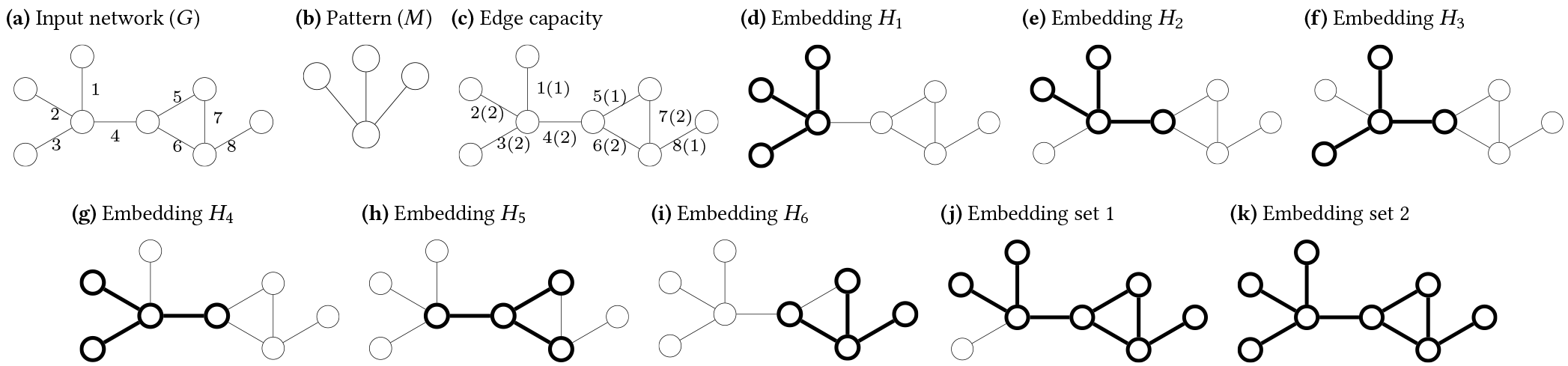
**(a) A network with eight nodes and eight edges. (b) A motif pattern. (c) A network with edge capacities. *x*(*y*) denotes the edge *x* has the capacity *y*. (d) - (i) six motif embeddings in the network. (j) an embedding set includes embeddings** *H*_2_, *H*_5_ **and** *H*_6_. **(k) an embedding set includes embeddings** *H*_1_, *H*_4_, *H*_5_ **and** *H*_6_.

Notice that both *F*_1_ and *F*_2_ measures make opposite, yet very strong assumptions regarding how the cells realize interactions. The former one assumes that the same interaction can participate in an arbitrary number of motif instances at the same time. The latter one limits the participation of an interaction to a single motif instance. Although, these two assumptions simplify the motif counting problem, they rarely reflect how cells operate interactions. Each interaction utilizes the molecules participating in that interaction. Thus the abundance of the interacting molecules makes it possible to include the corresponding edges appear in multiple (yet limited) number of motif instances. For example, depending on the concentration level of a metabolite, an enzymatic reaction may take place in one (low reaction/edge capacity) or many metabolic pathways (high edge capacity) [4, 9, 40]. We use the term *capacity* of an edge to denote the number of motif instances that edge can participate simultaneously. Thus, even if two cells have the same underlying biological network topology, they may yield different number of motifs of the same topology. For example, if we allow partial overlap of the motif *M* in the network *G* (see Figures 2b and 2c), we find four possible embeddings of motif *M* in *G* (Figures 2d, 2g, 2h and 2i). *A motif counting approach that ignores the edge capacities will lead to unrealistic motif counts, and wrong biological interpretations. Motif counting approaches that take edge capacities into account are needed*.

**Our contributions.** In this study, we build a new motif counting algorithm that allows partial overlap between different embeddings of a given motif on the target network. Briefly, given a target network, a motif topology, and a positive capacity value for each interaction in the target network, we count the maximum number of ways to place the motif on the target network, so that no edge appears in more motif embeddings than its capacity. Notice that the classical counting measures *F*_1_ and *F*_2_ are special instances of our measure [20, 36]. If we set the capacity of all edges to one, our motif counting problem reduces to non-overlapping motif counting with *F*_2_ measure. Similarly, if we set the edge capacities to infinity our problem reduces to motif counting using *F*_1_ measure.

We develop a novel motif counting method called *Partially Overlapping MOtif Counting* (*POMOC*), that computes the number of partially overlapping instances of a given motif in a given network. *POMOC* algorithm first finds all instances of a given motif *M* in the network *G*, then it chooses the motif instances that are guaranteed to exist without using any edge more than its capacity. For a given motif embedding, if the capacities of its edges are more than the number of embeddings those edges are part of, this embedding exists in our solution. Next, for the remaining motif instances, our algorithm randomly adds some embeddings whose edges are not more than the capacity constraints, into the resulting embedding set. It gradually expands the resulting set by taking one embedding out of the resulting set and inserting another two embeddings to the set.

Our experimental results on synthetic and real datasets demonstrate that our algorithm finds vastly different motif counts than *F*_1_ and *F*_2_ measures. Since biological interactions are resource limited, leading to varying edge capacities, we hypothesize that our method provides a more accurate approach to counting network motifs in biological networks than existing methods. Although, the *POMOC* method is slightly slower than motif counting with *F*_1_ and *F*_2_ measures, our method remains to be practical for all network sizes and motifs we test here. Our results on a *S. cerevisiae* transcriptional regulatory network suggest that oxidative stress is more disruptive to abundance and organization of network motifs than genetic mutations. Our analysis on the yeast network also suggests that our method can be used to find the key genes, which lead to topological and functional differences in biological networks under varying genetic backgrounds and growth conditions.

The rest of the paper is organized as follows. We present our algorithm in Section 2. We experimentally evaluate our method in Section 3 and provide a brief conclusion in Section 4.

## 2 METHOD

Here, we describe our method for counting partially overlapping motifs in networks. Section 2.1 provides the preliminaries needed to describe our method. Section 2.2 discusses our algorithm.

### 2.1 Preliminaries and problem definition

We denote a given biological network with graph *G* = (*V*, *E*,*c*), where *V* = {*ν*_1_, *ν*_2_,…, *ν*_*n*_} and *E* = {*e*_1_, *e*_2_,…, *e*_*m*_} represent the set of nodes (molecules) and the set of edges (interactions) among those nodes, respectively. The function *c*: *E* → ℤ^+^ shows the capacity of the edges. To simplify our notation, in the rest of this paper, ∀*e*_*i*_; ∈ *E* we use *c*_*i*_ to denote *c*(*e*_*i*_) (i.e., the capacity of the edge *e*_*i*_) and the vector *C* = (*c*_1_, *c*_2_,…, *c*_*m*_) to represent the capacity of all edges in *E* in sorted order of edge indices. For example, in Figure 2c, the value in the form *x*(*y*) denotes that edge *x* has capacity *y*.

Given a motif pattern *M*, we represent the *i*th embedding of *M* in *G* with *H*_*i*_ ⊆ *E* (i.e., *H*_i_ constitutes a subgraph of *G*, which is topologically isomorphic to *M*). We denote the set of all possible embeddings of *M* in *G* with 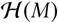. Given an edge *e*_*i*_ ∈ *E* and a subset 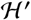 of 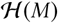, we denote the set of all embeddings in 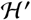 containing edge *e*_i_ with 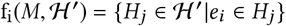. We denote the number of embeddings in the set 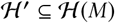 containing each interaction in *E* with the vector 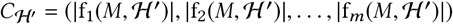. We say that 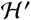 is *feasible* if no interaction appears in more embeddings in 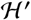 than its capacity that is 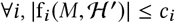. Figures 2b to 2i explain this on an example. Here, the capacity of the edges is *C* = (1, 2,2, 2,1, 2, 2,1). This network yields six embeddings (see Figures 2d to 2i). Consider the subset 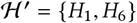. The capacity 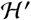 uses is 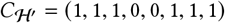, which is less than the imposed capacity constraints in *C*. Thus, 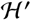 is feasible. The subset of embeddings {*H*_1_, *H*_2_}, however, is not feasible as this set contains edge *e*_1_ twice, which is more than its capacity (*c*_1_ = 1).

Next, we formally define the partially overlapping motif counting problem.

#### Definition 2.1.

(PARTIALLY OVERLAPPING MOTIF COUNTING). Consider a graph *G* = (*V*, *E*, *c*) conditioned with edge capacity constraints. Given a motif pattern *M*, partially overlapping motif counting problem seeks to find a largest feasible subset of the set of motif embeddings 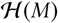.

Notice that Definition 2.1 provides a general formulation of motif counting problem. When the capacity constraints of each edge is set to infinity it reduces to a motif counting problem using *F*_1_ measure. When the capacity of all edges are set to one, it counts non-overlapping motifs (i.e., *F*_2_ measure). Next, we present the partially overlapping motif counting problem on an example.

*Example 2.1*. Different ways to select embeddings leads to different number of possible embeddings. Consider the network in Figure 2c with capacity constraints *C* = (1, 2,2, 2,1,2, 2,1). Embedding set 1 (see Figure 2j) includes embeddings *H*_2_, *H*_5_ and *H*_6_, and is feasible as its usage of the capacity is (1,1, 0, 2,1,2,1,1). Embedding set 2 (see Figure 2k) includes embeddings *H*_1_, *H*_4_, *H*_5_ and *H*_6_. This set is also feasible. The number of embeddings in this set is four. The partially overlapping motif count is thus four as this is the largest set of feasible embeddings.

Counting partially overlapping motifs is an NP complete problem for several reasons. First, it requires solving the subgraph isomorphism problem, which is NP-complete [10]. Furthermore, as we explain above the *F*_2_ count is a special instance of the partially overlapping motif counting problem as we can reduce the nonoverlapping motif counting problem to partially overlapping motif counting by setting the capacity of all edges to one. The Maximum Independent Set (MIS) problem, which is NP-complete [15], reduces to the non-overlapping motif counting problem [20, 30]. Thus the partially overlapping motif counting problem requires solving at least two NP-complete problems. In this paper, we develop a scalable method to tackle this problem using the local search strategy.

**Figure 3.**
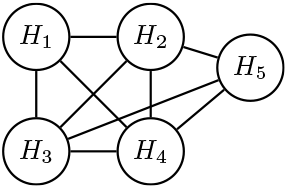
The overlap graph of the five embeddings in Figure 2d-2h.

### 2.2 Counting partial overlapping motifs

In this section, we discuss our *POMOC* algorithm. Algorithm 1 presents the pseudo-code of our method. Our algorithm takes a network *G* = (*V*, *E*, *c*) and a motif pattern *M* as input. Briefly, our algorithm has four main steps: (1) We locate all possible embeddings of *M* in *G* (line 1). At this step, we ignore the number of embeddings of *M* sharing each edge. (2) We determine the embeddings, which are guaranteed to exist in the final solution (lines 2-5). (3) We construct an initial, random yet feasible, solution by including a subset of the remaining embeddings in the set found in Step 2 (lines 5-6). (4) We iteratively improve this solution by replacing an embedding in the current solution with two or one new embeddings without violating feasibility of the solution (lines 7-11).

The first step of our algorithm is identical to computing the *F*_1_ count. This is a well studied problem in the literature. We use the method developed by Elhesha *et al* [12] for this step as it is one of the most recent and efficient methods. One can however replace this step with another method for *F*_1_ count without affecting the rest of our algorithm. Below, we explain Steps 2, 3, and 4 in detail.

*Step 2*. An embedding *H*_*r*_ is guaranteed to exist in the solution set if each edge of *H*_*r*_ has large enough capacity to realize all embeddings that have this edge. Formally, *H*_*r*_ exists in result set if 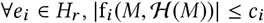. For example, in the network in Figure 2c, embedding *H*_6_ (Figure 2i) satisfies this criteria. We prove the correctness of this step at the end of this section.

*Step 3*. Once we identify the set of all embeddings, which are guaranteed to be in the final solution, we move them from the set 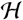 into solution set 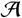. We then update the capacity constraints in the input graph *G* as follows. For each embedding 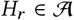, we reduce the capacities of all the edges *e*_i_ ∈ *H*_*r*_ by one as *H*_*r*_ is in the solution set. We then build a new graph, called the *overlap graph* for the remaining embeddings in 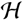. Each node in this graph corresponds to an embedding in 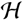. We include an edge between two nodes if their corresponding embeddings share at least one edge. Figure 3 depicts the overlap graph of the remaining five embeddings after moving embedding *H*_6_ from 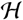 into the result set. Next, we generate a random feasible solution iteratively using the overlap graph as follows. At each iteration, we randomly pick a node *ν*_*r*_ from the overlap graph, and include the corresponding embedding (say *H*_*r*_) in the solution. We then reduce the capacity of all the edges of *H*_*r*_ in *G* by one. If the capacity of an edge drops to zero, it means that that edge cannot participate in any other embedding (say *H*_*s*_) without violating the feasibility of solution. If such an embedding *H*_*s*_ exists, its corresponding node in the overlap graph is a neighbor of *ν*_*r*_ as they share that edge in *G*. Thus, we remove these neighbors of *ν*_*r*_ in the overlap graph, which denote an embedding with an edge of zero capacity. We repeat these iterations to grow the random feasible solution set until the overlap graph becomes empty.

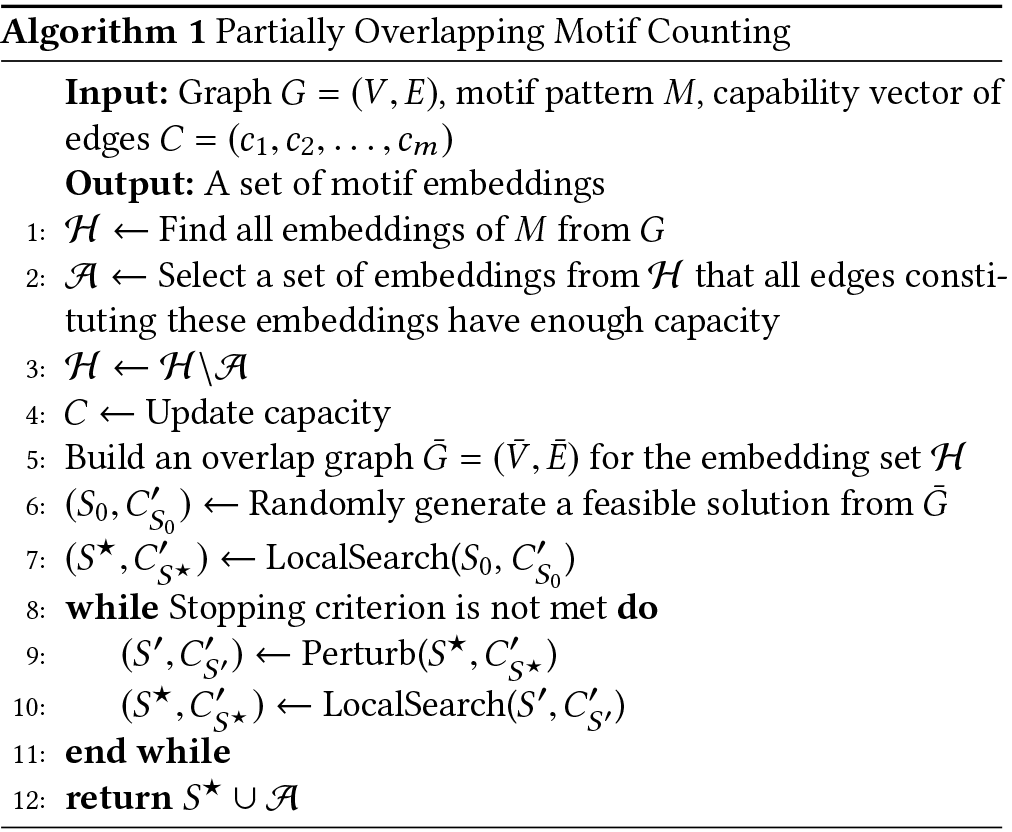

*Step 4*. So far, we have generated an initial feasible solution, which consists of those embeddings, which are guaranteed to be in the final result (Step 2) and those chosen randomly (Step 3). Next, we try to increase the size of this solution set iteratively by applying two strategies, namely *local search* and *perturbation*. We elaborate on these strategies next.

Local search iteratively increases the size of the solution set by replacing an embedding in the solution set with two other embeddings while maintaining the feasibility of the solution. For each embedding *H*_*i*_ in the solution, we study its corresponding node *v*_*i*_ in the overlap graph along with the set of neighboring nodes *v*_*i*_. We find all pairs of nodes (*ν*_*j*_, *ν*_*k*_) in the overlap graph, which satisfy all of the following two conditions.

i. *ν*_*j*_ and *ν*_*k*_ are neighbors of *ν*_*i*_.
ii. Replacing the embedding *H*_*i*_ corresponding to *ν*_*i*_ in the solution set with those of *H*_*j*_ and *H*_*k*_ corresponding to nodes *ν*_*j*_ and *ν*_*k*_ respectively does not violate the feasibility of the solution.

Once we identify all such node pairs, we randomly pick one and replace the embedding for node v; with those of *ν*_*j*_ and *ν*_*k*_ After each swap, we update the capacity of the edges contained in the embeddings *H*_*i*_, *H*_*j*_ and *H*_*k*_. Notice that each swap operation increases the solution set size by one. Also notice that the strategy described above may miss some nodes whose embeddings can be included in the solution set while maintaining the solution feasibility. This is possible, when the overlap graph has several components and there is no node of one component in the solution, the local search process will never consider this component. In order to take such nodes into consideration, after all possible swaps are performed, we find all non-solution nodes, which do not have neighbors among the solution nodes. We then insert the nodes in this set one by one to the solution if the solution remains feasible.

Similar to many local search algorithms, *POMOC* has the potential to get trapped in the local optimum. To escape the local optimum, before each local search, *POMOC* perturbs the current best solution to explore a slightly different search space. More specifically, *POMOC* first selects a node *ν*_*i*_ such that *H*_*i*_ is in the solution set and replaces it with an embedding *H*_*j*_ such that *ν*_*j*_ is a neighbor of *ν*_*i*_ in the overlap graph if the solution set remains feasible. As a result, the size of the solution set remains unchanged after the perturbation. For each node in the solution, *POMOC* does a Bernoulli trial with a probability value *p* to determine if a replacement is needed for this node. Given a solution *S*, *POMOC* does |S| × *p* replacements approximately. If a replacement is decided, *POMOC* finds all valid non-solution neighbors and randomly pick one to replace. Here, valid neighbors means the replacement of these nodes is consistent with the capacity constraints.

We repeat updating the solution by applying local search and perturbation until the size of the solution set does not improve for user supplied number of iterations. Finally, we prove the correctness of Step 2 of our algorithm in the following theorem.

#### Theorem 2.2.

Given a graph *G* = (*ν*, *E*, *c*) conditioned with edge capacity constraints and a motif pattern *M*, consider the set of all possible embeddings of *M*, 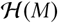. An embedding 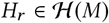 must be included by partially overlapping motif counting problem, if 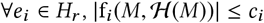.

**Proof.** We prove the theorem by contradiction. Assume that 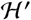 is the largest feasible subset of the set of motif embeddings 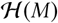, which does not have *H*_*r*_. That is, 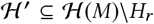. Now we construct a larger subset of embeddings 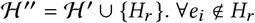, we have 
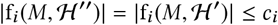

In addition, ∀ *e*_*i*_ ∈ *H*_*r*_, we have

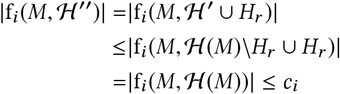

Thus, 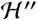 is also a feasible subset of embeddings, which yields a contradiction to the assumption that 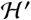 which does not include *H*_*r*_ is the largest feasible subset of 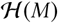. Thus, an embedding *H*_*r*_ must be included by partially overlapping motif counting problem if 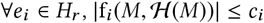

## 3 RESULTS

In this section, we experimentally evaluate performance of our method on synthetic and real datasets. We consider four motif topologies, which are commonly studied in the literature; namely bifan, biparallel, cascade and feed forward loop (see Figure 1). These motifs have been shown to be overrepresented in biological networks under the *F*_1_ count [23, 31]. We compare our method’s motif count and running time to two existing approaches: counting with *F*_1_ and *F*_2_ measures. First, we describe in detail the synthetic and real datasets used in our experiments. We then present the results.

### 3.1 Datasets

SYNTHETIC DATASET. We generate directed random networks to test robustness and scalability of our algorithm under three network parameters: network size (i.e., number of nodes), edge capacity, and topology model. We generate synthetic networks of varying sizes and edge capacities using three network topology models: Erdos-Renyi random graph (ER) [13], Watts-Strogatz small-world (WS) [38] and Barabasi-Albert preferential attachment (BA) [5]. We randomly assign the direction of each edge. We set a capacity value to each edge using two alternative approaches. In the first approach, we set the capacity of all edges to the same value; 1, 2 or 3. In the second approach, we randomly assign each edge capacity to a value between 1 and 3, thus different edges can have different capacities.

REAI DATASET. We use *S. cerevisiae* transcription regulatory network [11, 23]. This network contains 690 nodes and 1081 edges. We use the *S. cerevisiae* gene expression dataset, *GSE26169*, obtained from the GEO database to set capacities of interactions [7]. This dataset contains expression data under control and oxidative stress conditions in seven genetic backgrounds; wildtype, and Glr1, Gpx1, Gpx2, Grx1, Grx2 and Yap1 mutants leading to 14 different conditions (i.e., 2 × 7). We assign the capacity of each network edge using the capacity of the reactant gene. For each condition, we calculate the capacity of each gene as log(*e*_*ɡ*_)/*κ*, where *e*_*ɡ*_ and *κ* represent the expression level of gene *ɡ* and capacity constant, respectively. In our experiments, we use *κ* = 2.

IMPLEMENTATION AND SYSTEM DETAILS. We implement the *POMOC* algorithm in C++. We perform all the computational experiments on a Linux machine equipped with Intel core i7 processor 3.6 GHz CPU and 12GBs RAM.

### 3.2 Evaluation on synthetic networks

In this section, we compare *POMOC* to motif counting with *F*_1_ and *F*_2_ measures on synthetically generated networks for varying network size, network topology and edge capacity.

#### 3.2.1 Effects of network size

We generate random networks of varying sizes (200, 400, 800 and 1600 nodes) using ER, WS and BA topology models. We set the average degree of nodes to six, and the capacity for each edge to two. For each network size and model we generate 10 networks, and report the average motif count and running time. Figure 4 reports the results.

*Effects of network size on motif count (Figures 4a to 4c)*. We observe that motif count using partial overlaps significantly differs from that of *F*_1_ and *F*_2_ measures. Motif counting with *F*_2_ measure finds substantially lower motif counts than our method for each motif type in all network topologies for almost all motif types. In ER networks, only biparallel motif count shows observable difference between partial overlap and *F*_1_ measure. In WS networks, feed forward loop and biparallel motifs show the highest differences. In BA networks, all motif topologies show difference; the most significant variation is observed for the biparallel motif. The *POMOC* method and motif counting with *F*_1_ measure shows the most significant difference in BA networks. This difference can be explained as follows. ER and WS models generate synthetic networks whose nodes have similar degrees leading to uniform motif distributions in these network topologies. However, BA networks contain hub nodes with high degrees, which leads to non-uniform distribution of motifs to edges. Thus all motif embeddings can be realized in ER and WS networks but not in BA without violating our edge capacity constraints.

All three motif counting methods show the least motif count in synthetic networks generated under the ER model. Motif counts in WS and BA networks are comparable, but substantially higher than those in ER networks. This difference can be explained by the fact that ER model generates networks that are more disconnected compared to WS and BA models. Our results also show that regardless of the network model and counting measure the motif count varies dramatically across different motif topologies. More importantly, the motif count distribution exhibits significant variance across different random network models (see Figures 4a to 4c). While biparallel motif is observed most in ER and BA networks, feed forward loop is the most abundant motif in WS networks. Cascade motif is found least in ER and BA networks, but bifan is the least abundant motif in WS networks.

*Effects of network size on running time (Figures 4d to 4f)*. Our results demonstrate that counting motifs takes the most time with partial overlap constraint, and least time with *F*_1_ measure for all network sizes, topologies and motif types. This is not surprising since *F*_2_ count and partial overlap require solving the *F*_1_ count as the first step. Furthermore, since *F*_2_ count enforces identifying non-overlapping set of motifs, it eliminates the motif embeddings identified in *F*_1_ count more aggressively than the partial overlap. That said, we observe that the running time of our method is either very close to those of *F*_1_ and *F*_2_ measures or remains to be practical for all network sizes and motifs we test.

We observe that the running times for all three measures are greatly affected by the underlying network topology. Although all running times are less than 1 second for ER and WS networks, they go up to 10 seconds for BA networks. The second factor that influences the running time of our method is the motif topology. While our method’s running time for bifan motif is largest on ER and WS networks, that for biparallel motif is largest on BA networks. Cascade motif takes the shortest time in all three network topologies due to its smaller abundance and simple three node topology.

Three motif counting approaches show network topology dependent differences. In ER networks, only running times for feed forward loop and biparallel motifs show observable difference for the three measures. Similarly, in WS networks, running times for feed forward loop, biparallel and cascade motifs differ. The running time for our method differs from that of the *F*_2_ measure by a larger margin on BA networks for all motif types and network sizes. The most significant gap is observed for the biparallel motif. Finally, we would like to note that running time is independent of the motif count in ER and WS networks. However, there is a positive correlation between motif count and running time for BA networks; running time increases with increasing motif count.

**In summary**, our experiments suggest that motif distributions are heavily impacted by the network topology. In particular, BA networks show the most difference between *POMOC* method, and motif counting with *F*_1_ and *F*_2_ measures. Since biological networks often have similar topological characteristics as BA networks [6], we conjecture that our method will be crucial in determining the motif counts for real networks. On the other hand, our experiments also suggest that *POMOC* method is slightly slower than motif counting with *F*_1_ and *F*_2_ measures. While all three network topologies show running time differences for three motif counting approaches, the most significant difference is observed in BA networks. In all three network models, we observe that as network size increases running time increases linearly. Our method, however, scales to large networks (it runs in less than 10 seconds even for a network with 1600 nodes). Thus, our method is scalable to genome scale biological networks.

#### 3.2.2 Effects of capacity

Here, we compare the performance of our method to that of motif counting with *F*_1_ and *F*_2_ measures under varying edge capacity levels. We compare these three approaches using two metrics: motif count and running time. Similar to the previous section, we generate random networks with size 800 and average node degree 6 using ER, WS and BA network topology models. The capacity for each network edge is set using two distinct approaches. In the first approach we set capacity of all edges to 2 (*C*_2_) or 3 (*C*_3_). In the second approach, for each network edge, we randomly set capacity to an integer between 1 and 3 (*C*_*R*_). Note that motif counting with *F*_1_ and *F*_2_ measures correspond to setting the edge capacity to infinity (*C*_*INF*_) and one (*C*_1_), respectively. For each edge capacity and network topology model we generate 10 random networks, and report the average and two standard errors of motif count and running time. Figure 5 shows the results.

**Figure 4:**
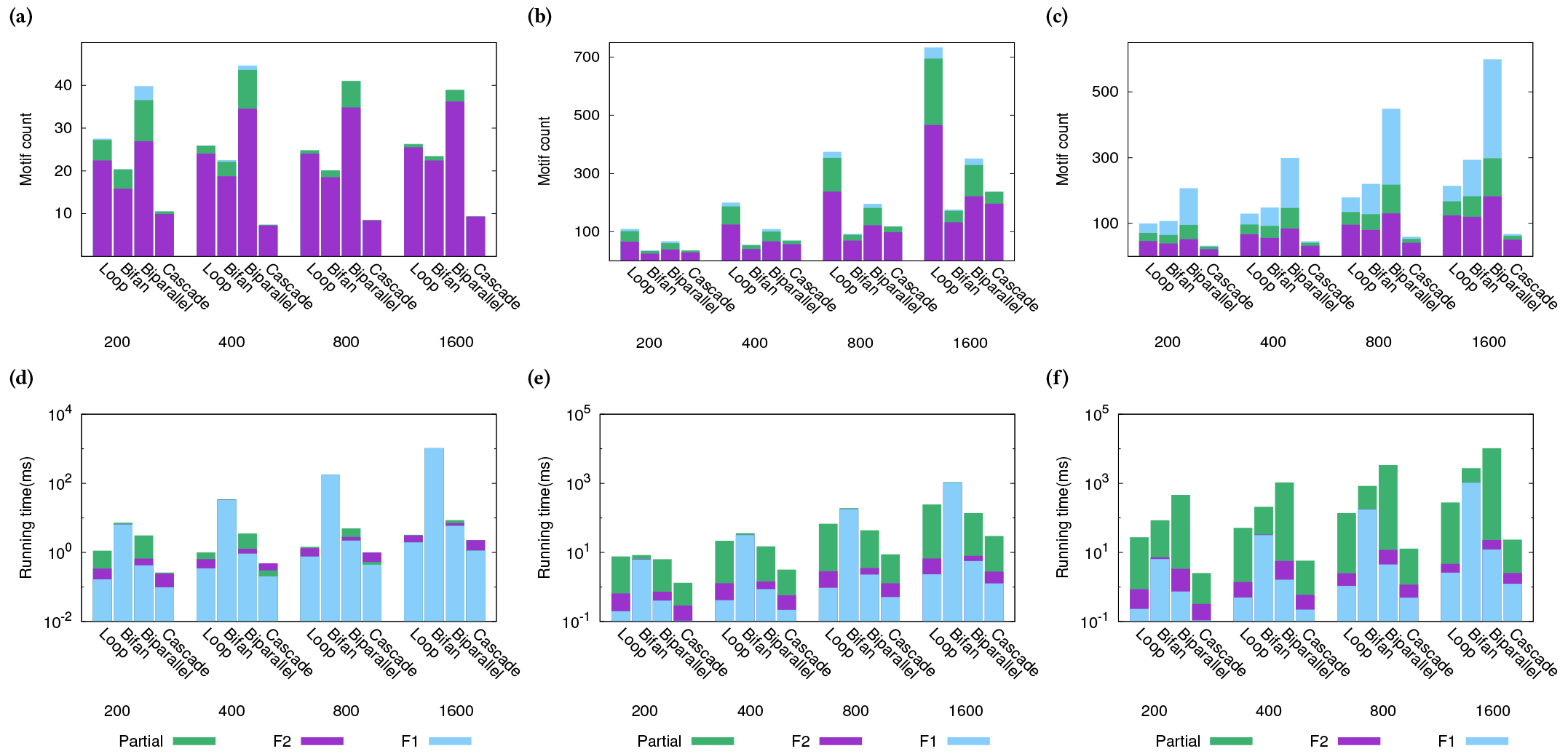
Motif count and running time on varying sizes of synthetic networks generated by (a, d) ER, (b, e) WS and (c, f) BA network topology models.

*Effects of capacity on motif count (Figures 5a to 5c)*. Our results suggest that the effects of edge capacity depends greatly on the network topology model. The motif counts for the WS and BA networks are substantially higher than that for the ER network. For the ER network, at all capacity levels, biparallel and cascade motifs are observed the most and the least, respectively. In WS networks, while feed forward loop is observed the most, bifan motif is observed the least in this network topology. In BA networks, biparallel and cascade motifs are observed the most and the least, respectively.

Next, we analyze how the motif counts change as edge capacity increases. We observe that as the capacity level increases the number of motifs found by the *POMOC* method tends to increase. This is not surprising as higher capacity values make it possible to have more motifs share the same edge without violating the capacity constraints. Motif counting with *F*_1_ (*C*_*INF*_) and *F*_2_ (*C*_1_) measures show the most and least number of motifs, respectively. Motif counting with varying edge capacities (*C*_*R*_) results in motif counts that are comparable to *C*_1_ and *C*_2_. Increasing edge capacity affects the motif counts in different ways for the three network topologies. In ER and WS networks, number of motifs are comparable between motif counting with *F*_1_ measure and partial motif counting with an edge capacity of 2 and 3, respectively. This suggests that for these two network topologies, a small edge capacity is enough to find all possible embeddings for each motif type. However, in BA networks, the motif counts are vastly different between motif counting with *F*_1_ measure and *POMOC* method. For this network topology, even an edge capacity of 3 is not enough to find all network motif embeddings. This observation can be explained by the fact that ER and WS models generate synthetic networks whose nodes have similar degrees. This leads to uniform motif distribution in these two network topologies with small number of motif overlaps. Thus, all motif embeddings could be found by using small edge capacities. However, the scale-free networks generated by BA model contain hub nodes with high degrees. The distribution of motifs to edges are non-uniform in these networks leading to large number of motif overlaps. Thus, all motif embeddings cannot be realized in BA networks without violating edge capacity constraints even with an edge capacity of 3-resulting in much lower motif counts as compared to the *F*_1_ measure.

*Effects of capacity on running time (Figures 5d to 5f)*. Our results demonstrate that our method is very fast for both fixed and variable capacity values. We observe that *POMOC* method’s running time for four motifs is affected by the network topology. The running time for all network motifs is less than one second in ER and WS networks. Although finding motifs in BA networks takes slightly longer, running time is still less than 5 seconds. Counting bifan and cascade motifs take the longest and the shortest time in ER and WS networks, respectively. Our algorithm shows different performance for BA networks. While bifan motif’s running time is highest in motif counting with *F*_1_ and *F*_2_ measures, biparallel motif count’s running time is highest for motif counting with edge capacity levels; *C*_2_, *C*_3_ and *C*_*R*_. For all capacity levels, cascade motif takes the shortest time in this network topology.

**Figure 5:**
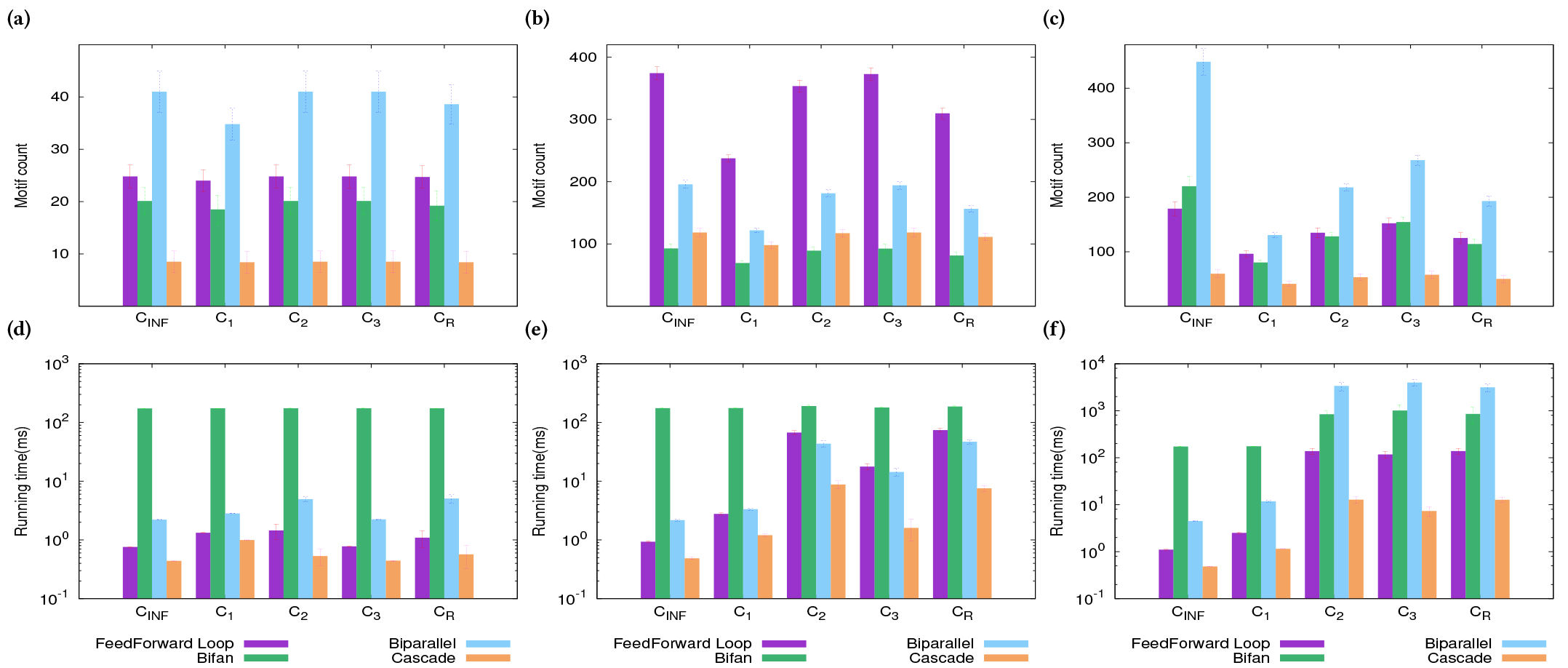
Motif count and running time on synthetic networks generated by (a, d) ER, (b, e) WS and (c, f) BA network models with varying edge capacity levels.

Running time for each motif changes similarly for ER and WS networks; in general motif counting with *C*_*R*_ and *C*_*INF*_ takes the longest and the shortest time, respectively. Interestingly, our method’s running time does not increase or decrease monotonically as edge capacity increases in these two network topologies. While running time increases from edge capacity of 1 to 2, the opposite behavior is observed as edge capacity increases from 2 to 3. Again, our algorithm shows slightly different behavior in BA networks. The running time increases as edge capacity increases from 1 to 2, but it is comparable for the capacity levels 2 and 3. In ER and WS networks, the running time does not correlate the motif counts. However, BA networks show positive association between motif count and running time.

**In summary**, our experiments suggest that BA networks will benefit from our partial motif counting method the most. Since, BA model generates scale-free networks that resemble the real biological networks, our method will potentially lead to more accurate motif counts and biological interpretations. On the other hand, our experiments also demonstrate that *POMOC* method is fast (less than 5 seconds) for a network of 800 nodes. Thus, we conjecture that that our method can scale to large real biological networks. Next, we test this conjecture by running our method on real networks.

### 3.3 Evaluation on real networks

In this section, we evaluate the performance of the *POMOC* method on a *S. cerevisiae* (budding yeast) transcriptional regulatory network. In our analysis, we focus on the effects of edge capacity to motif counting in seven genetic backgrounds (wild type, and Gpx1, Gpx2, Grx1, Grx2, Glr1 and Yap1 mutants) under two experimental conditions (normal and oxidative stress). As explained above, the *POMOC* method uses gene expression levels to calculate the network edge capacities. Thus, we use our method to count motifs of the yeast transcription network for 14 edge capacity vectors. Tables 1-2 and Figure 6 present our results. The running times for motif counting with *F*_1_ and *F*_2_ measures are less than 1 second in all real network experiments. *POMOC* method’s running time-in average 22 seconds-is comparable to those of *F*_1_ and *F*_2_ measures, and remains to be practical for real biological networks.

**Table 1:** Average and standard deviation of the motif counts for four motif topologies using the *POMOC* method and, motif counting with *F*_1_ and *F*_2_ measures.

|  | Feedforward | Bifan | Biparallel | Cascade |
| --- | --- | --- | --- | --- |
| $F_1$ | 73 | 2026 | 11 | 1 |
| $F_2$ | 24 | 137 | 5 | 1 |
| *POMOC* | $54 \pm 0.5$ | $426.4 \pm 7$ | $10 \pm 0$ | $1 \pm 0$ |

#### 3.3.1 Average motif counts

Here, we compare the average motif counts for four motif topologies over the 14 input networks using the *POMOC* method and, motif counting with *F*_1_ and *F*_2_ measures. Table 1 reports the mean and standard deviation of motif count for each counting approach. Note that *F*_1_ and *F*_2_ measures do not utilize edge capacity, and thus, their motif counts do not change under varying edge capacity vectors. Our results show that *POMOC* finds substantially different number of motifs than that with *F*_1_ and *F*_2_ measures. We observe that there is a massive variation in the abundance of different motif topologies from only one up to over two thousand. There is only one instance of cascade. Thus, this motif instance occurs in all three measures as it has no other instance to overlap with. More importantly, partially overlapping motif counting varies greatly in the interval defined by *F*_1_ and *F*_2_ measures for different motif topologies. For instance, for biparallel, it is closer to *F*_1_ measure, while it is closer to *F*_2_ measure for bifan, and equally distant to *F*_1_ and *F*_2_ measures for feed forward loop. This demonstrates that the amount of overlap among different embeddings of a motif depends not only on the network topology but also the motif topology. Similar to Milo *et al.* (2002) [23], we find that bifan and feed forward motifs are the most abundant motif types in this network. These two motif topologies also show small variation across the 14 networks for the *POMOC* method. Finally, our results show that the motif count distribution in yeast network differs substantially from that of all three synthetic network models we test (see Figures 4a to 4c). Biparallel motif is the most abundant in ER and BA networks, and feed forward is the most abundant in WS network. On the other hand, bifan is the most abundant motif in yeast network with a huge margin.

**Figure 6:**
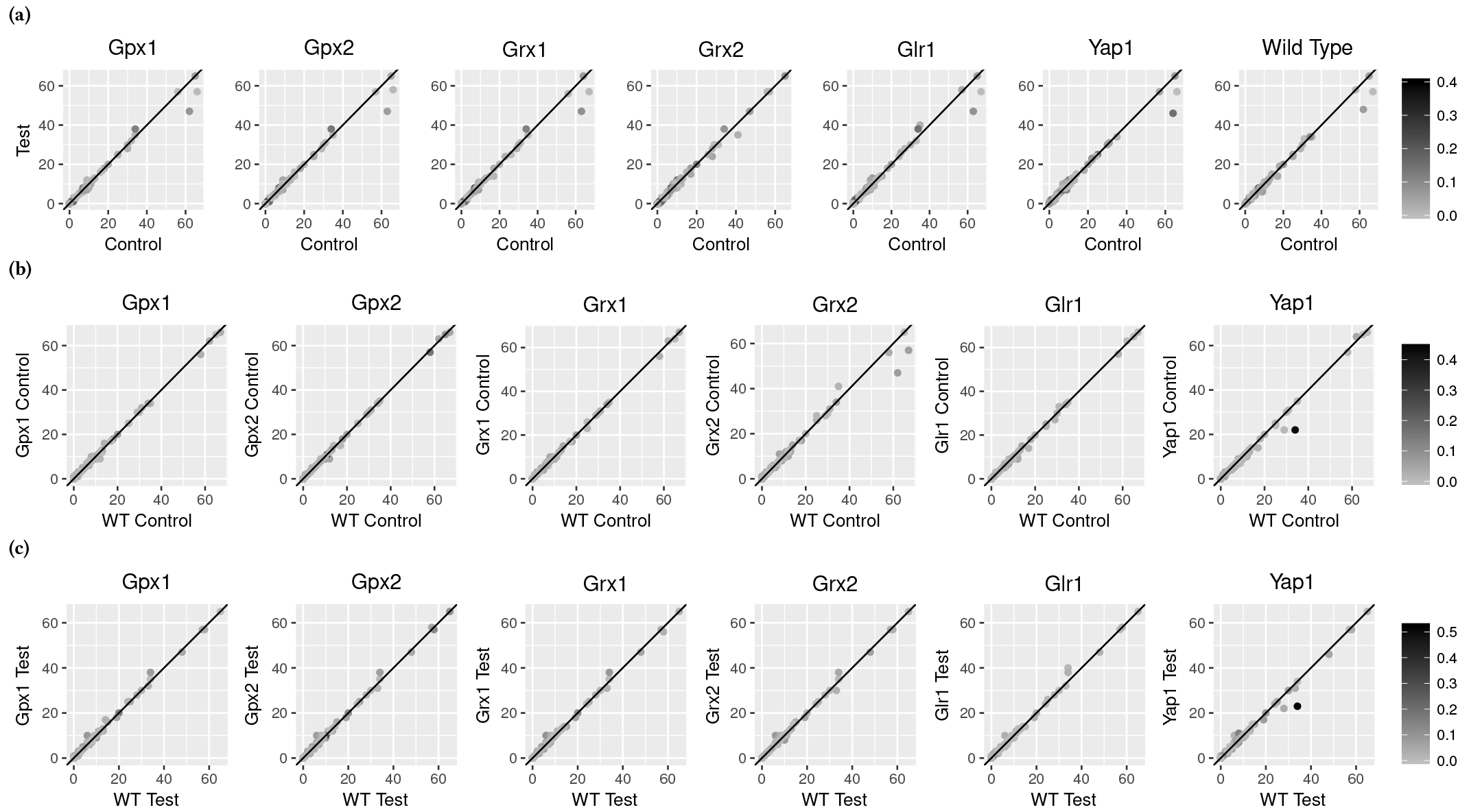
Motif counts for all reactant genes under control and oxidative stress conditions in seven genetic backgrounds (a), and in wild type and six genetic mutants under control (b) and oxidative stress (c) conditions.

**In summary**, our partial motif counting approach finds vastly different motif counts than *F*_1_ and *F*_2_ measures. Since real biological interactions are resource limited, leading to varying interaction (edge) capacities, we hypothesize that our method provides a more accurate approach to count network motifs, and to decipher dynamics of biological systems under varying conditions. Our method shows little variation in motif counts on the yeast transcription network (see the standard deviation values in Table 1), which suggests that variation in gene expression does not substantially change the edge capacity levels for the yeast network. That said, by focusing on the edges, which lead to such minor variation can reveal subtle differences between the impact of different cell states. Next, we analyze the association between motif count and edge capacity focusing on individual genes.

#### 3.3.2 Gene specific motif counts

Here, we take a closer look into the network contents and analyze the distribution of motif embeddings to individual genes. To do that, we first find the set of partially overlapping motif embeddings for each of the 14 genetic background and experimental condition combination. For each gene in each network, then, we count the number of motifs containing that gene. Figure 6 plots these counts for all reactant genes that we use to assign the edge capacities in the *S. cerevisiae* transcriptional regulatory network. In what follows, we discuss the motif count differences and effects of the edge capacity levels to motif counting in seven genetic backgrounds under control and oxidative stress conditions.

**Effects of experimental condition to motif count**. Our results demonstrate that most of the genes exhibit similar motif counts under varying genetic backgrounds and growth conditions. We observe more genes away from the *y* = *x* line in control *vs*. oxidative stress conditions (Figure 6a) than wild type *vs*. mutant genetic backgrounds (Figures 6b and 6c). This suggests that oxidative stress is more disruptive to the organization and abundance of motifs in yeast network than mutations of individual genes.

To understand our results more clearly, from now on, we focus on the analysis of the six reactant genes; Abf1, Dal80, Msn2, Msn4, Skn7 and Yap1. Among these, Msn2, Msn4, Skn7 and Yap1 play key roles in controlling the transcription of oxidative stress response genes [22, 24, 25, 34, 35]. Dal80 is implicated in nitrogen depletion and amino acid starvation stress responses [26, 39]. Similarly, Abf1 is involved in nutritional stress response [14]. Table 2 reports how changes in the edge capacity levels affect the number of motifs these six genes participate. For this analysis, first we divide the edge capacity and motif count variations to two groups: significant or non-significant. We designate any variation that is more than 10% as significant. We then divide all reactant genes to four groups (00, 01, 10 and 11). Here, the first digit represents whether observed variations in edge capacity is significant (1) or not (0), respectively. Second digit denotes whether the deviation in motif count is significant.

**Table 2:** Motif count variation for Abf1, Dal80, Msn2, Msn4, Skn7 and Yap1 under control and oxidative stress conditions in seven genetic backgrounds (a), and in wild type (WT) and six genetic mutants under control (b) and oxidative stress (c) conditions. The first and second digits represent whether the observed difference in edge capacity and motif count is significant (1) or not (0).

|  |  | Gpx1 | Gpx2 | Grx1 | Grx2 | Glr1 | Yap1 | WT |
| --- | --- | --- | --- | --- | --- | --- | --- | --- |
| (a) | Abf1 | 00 | 00 | 00 | 01 | 00 | 01 | 01 |
|  | Dal80 | 00 | 00 | 00 | 10 | 00 | 00 | 00 |
|  | Msn2 | 01 | 01 | 01 | 00 | 01 | 01 | 01 |
|  | Msn4 | 01 | 01 | 01 | 00 | 01 | 11 | 01 |
|  | Skn7 | 00 | 00 | 00 | 00 | 00 | 00 | 00 |
|  | Yap1 | 11 | 11 | 11 | 01 | 11 | 00 | 00 |
|  |  | Gpx1 | Gpx2 | Grx1 | Grx2 | Glr1 | Yap1 |  |
| (b) | Abf1 | 00 | 00 | 00 | 00 | 00 | 01 |  |
|  | Dal80 | 00 | 00 | 00 | 00 | 00 | 00 |  |
|  | Msn2 | 00 | 00 | 00 | 01 | 00 | 00 |  |
|  | Msn4 | 00 | 00 | 00 | 01 | 00 | 00 |  |
|  | Skn7 | 00 | 00 | 00 | 00 | 00 | 01 |  |
|  | Yap1 | 00 | 00 | 00 | 00 | 00 | 11 |  |
|  |  | Gpx1 | Gpx2 | Grx1 | Grx2 | Glr1 | Yap1 |  |
| (c) | Abf1 | 01 | 01 | 11 | 01 | 01 | 01 |  |
|  | Dal80 | 10 | 10 | 10 | 00 | 00 | 00 |  |
|  | Msn2 | 00 | 00 | 00 | 00 | 00 | 00 |  |
|  | Msn4 | 00 | 00 | 00 | 00 | 00 | 00 |  |
|  | Skn7 | 00 | 00 | 00 | 00 | 00 | 01 |  |
|  | Yap1 | 01 | 01 | 01 | 01 | 01 | 11 |  |

First, we focus on motif count variation in seven genetic backgrounds under control and oxidative stress conditions (Figure 6a and second digits in Table 2a). Our analysis shows that while most of the reactant genes including Dal80 and Skn7 have similar motif counts, some genes such as Msn2, Msn4 and Yap1 show substantially different motif counts. We observe that motif count differences are highly dependent on genetic background: Msn2 and Msn4 show different motif counts in all genetic backgrounds except Grx2 mutant; Yap1 shows different motif counts only in Gpx1, Gpx2, Grx1, Grx2 and Glr1 mutants; Abf1 shows different motif counts only in wild type and, Grx2 and Yap1 mutants.

Motif counts show small variation between wild type and mutant conditions under control and oxidative stress conditions (Figure 6b-6c and and second digits in Table 2b-2c). Our results suggest that genetic mutants show similar gene expression levels to wild type, which leads to comparable edge capacity levels and motif counts. Under control conditions: Msn2 and Msn4 show motif count variation in Grx2 mutant; Yap1, Skn7 and Abf1 show motif count variation in Yap1 mutant. We observe slightly different behavior under oxidative stress conditions: Yap1 and Abf1 show motif count variation in all genetic backgrounds; Skn7 shows motif count variation in Yap1 mutant. Msn2, Msn4 and Dal80 show similar motif counts under oxidative stress in all genetic backgrounds.

**Effects of edge capacity to motif count**. Here, we analyze how an individual gene’s edge capacities affect the number of motifs it participates. Figure 6 shows that while genes with highly different edge capacity (darker circles) might have no motif count variation (circles on the *y* = *x* line); genes with similar edge capacity (lighter circles) might have motif count variation (circles off the *y* = *x* line). This implies that variation in edge capacity levels is not a sufficient condition for alterations in motif count. Also small variations in edge capacities may lead to substantial changes in motif counts. We will further analyze these observations on six stress response genes. The two digits in Table 2 reports the dependency between edge capacity and motif counts.

First, we report how oxidative stress affects motif counts. We observe that although oxidative stress leads to edge capacity variation for many genes, most of the genes show similar motif counts under control and oxidative stress conditions in seven genetic backgrounds. Msn2 and Msn4 show small edge capacity variation, but high motif variation in all genetic backgrounds except Grx2 mutant (Table 2a). Abf1 shows similar edge capacity in all genetic backgrounds, however, it shows different motif counts in wild type and, Grx2 and Yap1 mutants. In Grx2 mutant, Dal80 shows edge capacity variation, but similar motif counts. For Yap1 we see a more complex story. It shows edge capacity variation in Gpx1, Gpx2, Grx1 and Glr1 mutants, which leads to different motif counts. However, while Yap1 shows similar edge capacity levels in wild type, and Grx2 and Yap1 mutants, it has different motif counts in Grx2 mutant.

In almost all genetic backgrounds, majority of the genes do not show edge capacity variation between wild type and mutant genetic backgrounds under control condition (Table 2b). Yap1 shows edge capacity and motif count variation in Yap1 mutant. Msn2 and Msn4 show similar edge capacity but different motif counts in Grx2 mutant compared with wild type. Similarly, Abf1 and Skn7 show motif count variation in Yap1 mutant. Oxidative stress leads to slightly more edge capacity variation compared to control (Table 2c). Yap1 and Abf1 show edge capacity and motif count variation in Yap1 and Grx1 mutants compared with wild type, respectively. Dal80 shows edge capacity variation in Gpx1, Gpx2 and Grx1 mutants, however, it has similar motif counts. Despite lack of edge capacity variation, Yap1 shows different motif counts between wild type and, Gpx1 Gpx2, Grx1, Grx2 and Glr1 mutants. Similarly, Abf1 does not show edge capacity variation in Gpx1, Gpx2, Grx2, Glr1 and Yap1 mutants, but, it has different motif counts. Finally, Skn7 shows motif count variation in Yap1 mutant although having similar edge capacity.

**In summary**, partial motif counting approach finds substantially different motif counts than that of *F*_1_ and *F*_2_ measures. However, the changes in edge capacity is not enough to explain the observed motif count differences for individual genes. While some genes show different motif counts due to edge capacity changes, most genes have the same motif count. There are also genes that show similar edge capacity levels but different motif counts. For these genes, the changes in motif counts could be possibly explained by the edge capacity changes of its neighboring genes.

## 4 CONCLUSIONS

Motif counting in biological networks is an important tool to decipher the topology of biological networks and its function. Existing motif counting approaches either count all or non-overlapping instances of a given motif. This results in either over-or underestimation of the motif counts, since biological reactions are constrained by many factors such as concentration levels of reactant molecules. In this paper, we presented a novel motif counting method, *POMOC*, based on edge capacities of a given network. Our experiments on both synthetic and real networks demonstrate that motif count using our method significantly differs from the existing motif counting approaches, and our approach extends to large-scale biological networks in practical time. Application of our method to the *S. cerevisiae* transcription regulatory networks demonstrates that the *POMOC* method reveals topological differences of biological networks under different genetic backgrounds and experimental conditions.

